# *Alternaria atra* from distinct ecological roles share functional genomic repertoires

**DOI:** 10.64898/2026.02.26.708203

**Authors:** Tamara Schmey, Karen Bahar, Christopher Tominello-Ramirez, German Sepúlveda Chavera, Remco Stam

**Affiliations:** Institute of Phytopathology, Department of Phytopathology and Plant Protection, Christian-Albrechts-Universität zu Kiel, Kiel, Germany; Departmento de Recursos Ambientales, Facultad de Ciencias Agronómicas, Universidad de Tarapacá, Arica, Chile

**Keywords:** Comparative genomics, Endophytism, Pathogenicity, Effectors, CAZymes, Lifestyle plasticity

## Abstract

Fungi, particularly ascomycetes, exhibit diverse ecological lifestyles, including endophytism, pathogenicity, and saprotrophy. Species of the genus *Alternaria* are taxonomically and ecologically diverse, yet the genomic determinants underlying different lifestyles remain poorly understood.

Here, we investigate lifestyle-associated genomic variation in *Alternaria atra* using two newly collected isolates obtained as plant endophytes. We confirm their taxonomic identity and generate draft genome assemblies for both isolates. We assess their phenotypic behaviour under laboratory conditions and examine their genomic features alongside those of a previously published *A. atra* isolate described as pathogenic.

Despite differing isolation histories, the endophytic and pathogenic isolates exhibit similar behaviour under laboratory conditions and possess highly comparable genomic repertoires, including predicted effector proteins, carbohydrate-active enzymes, and biosynthetic gene clusters. We detect no clear genomic signatures distinguishing endophytic and pathogenic origins or lifestyles.

These findings suggest that *A. atra* harbours a shared genomic repertoire compatible with multiple ecological strategies, supporting a model of lifestyle plasticity rather than fixed genomic specialization. Our results add to growing evidence that genome content alone does not reliably predict ecological roles in ascomycete fungi.

## Introduction

Fungi are among the most diverse and ecologically versatile eukaryotes, exhibiting lifestyles that range from free-living decomposers to obligate symbionts and pathogens. Despite their ecological importance, the genomic determinants underlying these lifestyles remain incompletely characterized, especially in non-model taxa. Genome sequencing and comparative analysis provide a means to connect fungal ecological strategies with their underlying genetic architecture.

Fungal lifestyles are defined by their modes of nutrition and interactions with hosts or substrates. Endophytic fungi colonize living plant tissues without causing visible disease symptoms (Schulz and Boyle 2005) and can be commensal or mutualistic, sometimes providing benefits like increased stress tolerance (Zanne et al. 2020). In contrast, pathogens cause disease when host defences are overwhelmed (Kogel et al. 2006). Endophytism and pathogenicity are often not discrete states but lie along a continuum, shaped by the balance between fungal virulence and host response (Schulz and Boyle 2005; Kogel et al. 2006).

Beyond host-associated lifestyles, saprotrophic fungi obtain nutrients by decomposing dead organic matter. Some fungi are capable of transitioning between saprotrophic and host-associated modes depending on environmental conditions and life-history stage (Schulz and Boyle 2005).

Some fungal genera show consistent lifestyles, for instance, all species in *Russula* are ectomycorrhizal and all species in *Puccinia* are plant pathogens (Zanne et al. 2020). However, across genera and species, isolates can exhibit diverse lifestyles, and even individual fungi can shift their mode of nutrition over time. An example for such shifts would be a forest fungus that can switch between saprotrophy and parasitism (Olson et al. 2012) or an endophytic population of *Verticillium dahliae* that evolved from a sympatric pathogenic population (Wheeler et al. 2019). Another example would be fusarioid fungi that transition between endophytism, saprotrophy, and pathogenicity (Ulaszewski et al. 2025). Using comparative genomics, Hill et al. showed that frequent lifestyle transitions in fusarioid fungi are poorly explained by canonical genomic predictors, underscoring the challenge of linking genome content to ecological function (Hill et al. 2022). Thomas et al. propose that some fungal endophytes use endophytism to enhance dispersal to woody substrates and persist through environmental stress, a concept formalized as the Foraging Ascomycete hypothesis (Thomas et al. 2020).

The genus *Alternaria* (Pleosporaceae, Ascomyctoa) is globally distributed (Lawrence et al. 2016) and ecologically versatile, as it can adopt different lifestyles (Thomma 2003). According to Thomma, most *Alternaria* species are saprotrophs in soil or decaying plant tissue (Thomma 2003).

Members of several *Alternaria* spp. can be endophytes. One example for endophytic growth is *Alternaria oxytropis* in locoweed (He et al. 2019). Zhang et al. 2025 found more *A. oxytropis* endophytes in regions with extreme winter conditions, leading them to speculate that the endophytes might provide an advantage in harsh environmental contexts (Zhang et al. 2025). *Alternaria alternata* can lead endophytic and pathogenic lifestyles, as reviewed in DeMers 2022. Aside from specificity conferred by host specific toxins that exist in some specific lineages, such as those infecting tangerine or strawberry, *A. alternata* lineages do not seem constrained to either endophytic or parasitic lifestyles, and likely all possess the capability for endophytism and mild pathogenicity (DeMers 2022).

Many species of *Alternaria* are economically relevant pathogens and cause yield losses on a wide range of important crops, including major food crops such as potato and tomato (Thomma 2003; Schmey et al. 2024). Furthermore, *Alternaria* spp. cause food spoilage at the postharvest stage. Alternaria have been isolated from different food products and can accumulate toxic metabolites (Patriarca 2016).

The genus *Ulocladium* was synonymized with the genus *Alternaria* and the species *Ulocladium atrum* became *Alternaria atra* (Woudenberg et al. 2013). It causes foliar diseases on potato plants (Nasr-Esfahani et al. 2021; Esfahani 2018). Despite being a pathogen on important crops, there are also studies about its suitability as biocontrol agent, for example against *Botrytis cinerea* (Elead et al. 1994). There is also a report of *A. atra* as a post-harvest rot, as a genome sequence was published from *A. atra* that was obtained from a supermarket Red Delicious apple after incubation (Gebru et al. 2020). The other genome sequence currently available is from an *A. atra* isolate that was collected from an infection lesion on a wild tomato relative (*Solanum chilense*) leaf in Chile (Bonthala et al. 2021). With only two full genomes available, *A. atra* is relatively understudied compared some other *Alternaria* species. For investigations into the genomic underpinnings of *Alternaria* lifestyles, additional genomes of this species might help fill knowledge gaps and enhance our understanding of its pathogenicity and ecological adaptability. Even the mating system of *A. atra* remains unresolved. Gannibal et al. 2024 reported *A. atra* as heterothallic, with isolates carrying either the MAT1-1-1 or MAT1-2-1 idiomorph (Gannibal and Gomzhina 2024). In contrast, Geng et al. 2014 had found both idiomorphs within the same isolate, suggesting homothallism (Geng et al. 2014).

*Tillandsia landbeckii* (Bromeliaceae) occurs in the driest regions of the Atacama Desert in Chile (Hakobyan et al. 2023) and is the dominant species forming *Tillandsia* lomas (Pinto et al. 2006). In such extreme habitats, drastic selection pressures are at play, making evolutionary studies of these plants and their endophytes particularly interesting. *T. landbeckii* hosts specific bacterial communities, providing refugia for microbial life in hyperarid desert environments (Hakobyan et al. 2023). In other habitats, researchers have identified fungal endophytes in different *Tillandsia* species (Tellez et al. 2020; Unterseher et al. 2013). Another plant species endemic to the Atacama Desert, *Malesherbia auristipulata*, also harbours endophytic *Alternaria* and *Fusarium* spp. (Sepúlveda Chavera et al. 2024).

In this study, we investigate the genomic underpinnings of *Alternaria atra*’s trophic modes and lifestyles. We describe the collection and whole genome sequencing of two isolates from symptomless *Tillandsia landbeckii* plants, identify them as *A. atra* through phylogenetic analysis, and characterize their growth patterns. We then compare these endophytic isolates to the pathogenic *A. atra* isolate CS162 under laboratory conditions. Using the genomes, we explore the genetic factors driving lifestyle adaptations by analysing effectors, biosynthetic gene clusters (BGCs), and carbohydrate-active enzymes (CAZymes).

## Methods

### Biological materials

The two new fungal isolates, T001 and T003, were collected on 17 January 2017 from asymptomatic *Tillandsia landbeckii* plants in outlying arid areas of the Atacama Desert near Arica. Due to the absence of visible symptoms on the host plants, both isolates are described as endophytic fungi. Purification and storage of the isolates followed the method described in (Schmey et al. 2023).

For laboratory experiments and comparative genomics, we analysed T001 and T003 alongside the previously published *Alternaria atra* isolate CS162 (Bonthala et al. 2021). The isolate CS162 represents a pathogen from a leaf lesion on a *Solanum chilense* plant near San Pedro de Atacama. The distance between the collection locations of the two new isolates and the collection location of CS162 is 501 km.

### Morphology

For morphological identification, we grew the isolates on SNA medium (Schmey et al. 2023) and took pictures of the conidia under the light microscope. Additionally, we took photos of the colony morphology after 7 days of growth on SNA medium.

### Sequencing

We isolated high-molecular-weight DNA following the phenol/chloroform protocol of Einspanier et al. 2022 (Einspanier et al. 2022). The LMU Gene Center (Munich, Germany) prepared the libraries and carried out sequencing using the Ligation Sequencing Kit SQK-LSK109 and the Native Barcoding Expansion kits (EXP-NBD104/114) on PromethION R11.1 and R9.4.1 flow cells. We sequenced the samples alongside additional, unrelated samples on the same flow cells and subsequently pooled the data from three such multiplexed runs. We performed basecalling with Guppy v5.0.13 (ONT, Oxford, UK) using the SUP (super-accurate) model. We evaluated sequencing output with NanoComp v1.23.1 (Coster and Rademakers 2023) and filtered reads with NanoFilt v2.8.1 (Coster et al. 2018), retaining only reads with a mean quality score above Q10.

### Assembly and annotation

For de novo genome assembly, we used Flye v2.9.5-b1801 (Kolmogorov et al. 2019) and subsequently polished using Medaka v2.0.1 (https://github.com/nanoporetech/medaka). We used QUAST v5.0.2 (Mikheenko et al. 2018) and BUSCO v5.8.2 (Manni et al. 2021) for the assembly evaluation. The annotation with funannotate v1.8.17 (Palmer and Stajich 2020) followed the same steps as Schmey et al 2025 (Schmey et al. 2025). We performed the annotation on the two newly assembled genomes and, to allow comparability, on the downloaded assemblies of CS162 (NCBI accession GCA_907166805.1) and MOD1FUNGI7 (NCBI accession GCA_004634305.1). The funannotate compare functionality helped compare the four genomes and their annotations.

### Mating types

We performed in silico PCR to screen the four genome assemblies for mating-type (MAT) genes using the four primer pairs described in (Gannibal and Gomzhina 2024). Two primer pairs target MAT1-1-1 (from (Gannibal and Kazartsev 2013; Geng et al. 2014)) and two target MAT1-2-1 (from (Berbee et al. 2003; Geng et al. 2014)). We conducted the in silico PCR with SeqKit v2.8.2 (Shen et al. 2024), allowing three mismatches and later four. All in silico PCR products were subjected to online BLAST searches against the NCBI nucleotide database (https://blast.ncbi.nlm.nih.gov/Blast.cgi) to confirm their identity.

We used BLASTn from BLAST+ v2.13.0+ (Altschul et al. 1990) to check for the presence of the two mating-type idiomorphs in all four assemblies. The query sequences for type MAT1-1-1 (accession HQ906703.1) and type MAT1-2-1 (accession HQ906727.1) came from the homothallic isolate *Alternaria atra* (formerly *Ulocladium atrum*) strain CBS 195.67 in the NCBI Nucleotide database.

### Phylogeny

To assess relationships among the sampled isolates, we applied two complementary tree-based approaches. First, we downloaded reference genomes from NCBI (accessions listed in figure 2) and inferred a cladogram using k-mer–based clustering in Mashtree v1.4.6 (Katz et al. 2019) with 1000 bootstrap replicates. We visualized the resulting cladogram and rooted it to *Stemphylium lycopersici* in iTOL (https://itol.embl.de). Second, we reconstructed a phylogeny based on barcode marker loci commonly used for *Alternaria* section *Ulocladioides*, namely *alt a 1, gapdh, tef1* and *rpb2*. We extracted marker sequences from genome assemblies using SeqKit and primer sequences previously employed by (Gannibal and Gomzhina 2024). We combined these sequences with corresponding reference sequences from (Gannibal and Gomzhina 2024) into locus-specific FASTA files. We used TrEase v2.0 (Mishra et al. 2023) with default settings to obtain cropped alignments. Then we concatenated the barcode marker alignments using Galaxy (https://usegalaxy.org) with the Fasta Concatenate by Species tool (https://toolshed.g2.bx.psu.edu/repos/devteam/fasta_concatenate_by_species/fasta_concatenate0/0.0.1). Finally, we used the concatenated alignment as input for TrEase to generate a RAxML phylogeny, which we visualized and rooted to *A. chartarum* in iTOL.

### Lifestyle under laboratory conditions

To confirm the endophytic and pathogenic lifestyles of the isolates under laboratory conditions, we conducted detached-leaf assays. We obtained *Tillandsia landbeckii* plants (Plantabrutt, Spain) and a *Solanum chilense* plant from accession LA4117A (TGRC, USA). For surface sterilization, we placed detached leaves in beakers with autoclaved, distilled water, then consecutively in 70% ethanol (EtOH) for 1 min, then 2% sodium hypochlorite (NaOCl) for 5 min, and finally in two beakers of autoclaved, distilled water. Afterwards, we placed the leaves on wet Whatman paper in glass dishes under a sterile bench. We cultured all fungal isolates on SNA medium (Einspanier et al. 2022) to promote sporulation. For the assay, we used the endophytic isolates T001 and T003, the known *S. chilense* pathogen CS162, and *Alternaria alternata* CS046 (from the collection of (Schmey et al. 2023)) as a positive infection control, alongside sterile water as a negative control. We prepared conidial suspensions by scraping conidia from the plates with glass slides, placing them in sterile water, and adjusting their concentration to 50,000 conidia per mL using a Thoma chamber. Afterwards, we added a drop of Tween to the suspensions and inoculated the leaves at the leaf base with 10 μL of the suspension. We monitored lesion development at approximately 21°C throughout the assay and photographed the leaves at 6 and 10 days post inoculation (dpi) to document symptom progression.

Following Koch’s postulates, we isolated the fungi from infected *T. landbeckii* and *S. chilense* leaves by transferring fungal material onto fresh SNA plates. For *T. landbeckii*, we dipped the lesion directly onto the medium, and for *S. chilense*, we scraped fungal tissue with sterile tweezers. After 10 days of incubation at approximately 21 °C, we identified the isolates using a light microscope.

### Comparative genomics for lifestyle indicators

The results from funannotate annotation include predictions of which of the proteins are secreted. We used these secreted proteins as input for effector prediction with effectorP-3.0 (Sperschneider and Dodds 2022) and plotted the results as stacked bar plot using RStudio.

During annotation, we ran antiSMASH v7.1.0 (Blin et al. 2021) and integrated the BGC results into the annotations. Here, we cluster the antiSMASH results using BiG-SCAPE v2 (Navarro-Muñoz et al. 2020) and visualize the presence absence matrices of gene cluster families.

Functional genome annotation with funannotate includes the annotation of CAZymes. Afterwards, we use the funannotate compare function to compare the CAZyme annotations between the four isolates. Then we re-plotted the CAZyme summary using Rstudio. We examined the abundances of the CAZyme families GH10, GH5, and AA9 in the funannotate compare output, as previous work linked these families to endophytism (Mesny et al. 2021; Yuan et al. 2022).

### Predictions of trophic modes using CATAStrophy

We used CATAStrophy v0.1.0 (Hane et al. 2019) to predict the trophic lifestyles of four isolates by running the protein files produced by our Funannotate annotations. CATAStrophy classifies filamentous plant pathogens by quantifying their CAZyme repertoires, projecting these frequencies into a PCA space trained on curated reference proteomes, and computing relative centroid distances (RCDs) to established trophic lifestyle classes. It assigns class membership by identifying the nearest class centroid and generating multi-label RCD scores that reflect each proteome’s similarity to all classes. We report the best prediction with an RCD score of 1 and provide ancillary table columns listing any additional classifications with RCD scores above 0.8.

## Results

### Morphology

Microscopic examination (figure 1) showed short, unbranched conidiophores with sympodial conidiogenesis. Conidia were solitary, dark brown to black, polytretic, and lacked beaks. They were obovoid, slightly larger than 20 μm, with transverse septation and a conspicuously verrucose surface. These features are consistent with section *Ulocladioides* as described in Woudenberg 2013 (Woudenberg et al. 2013).

**Figure 1.**
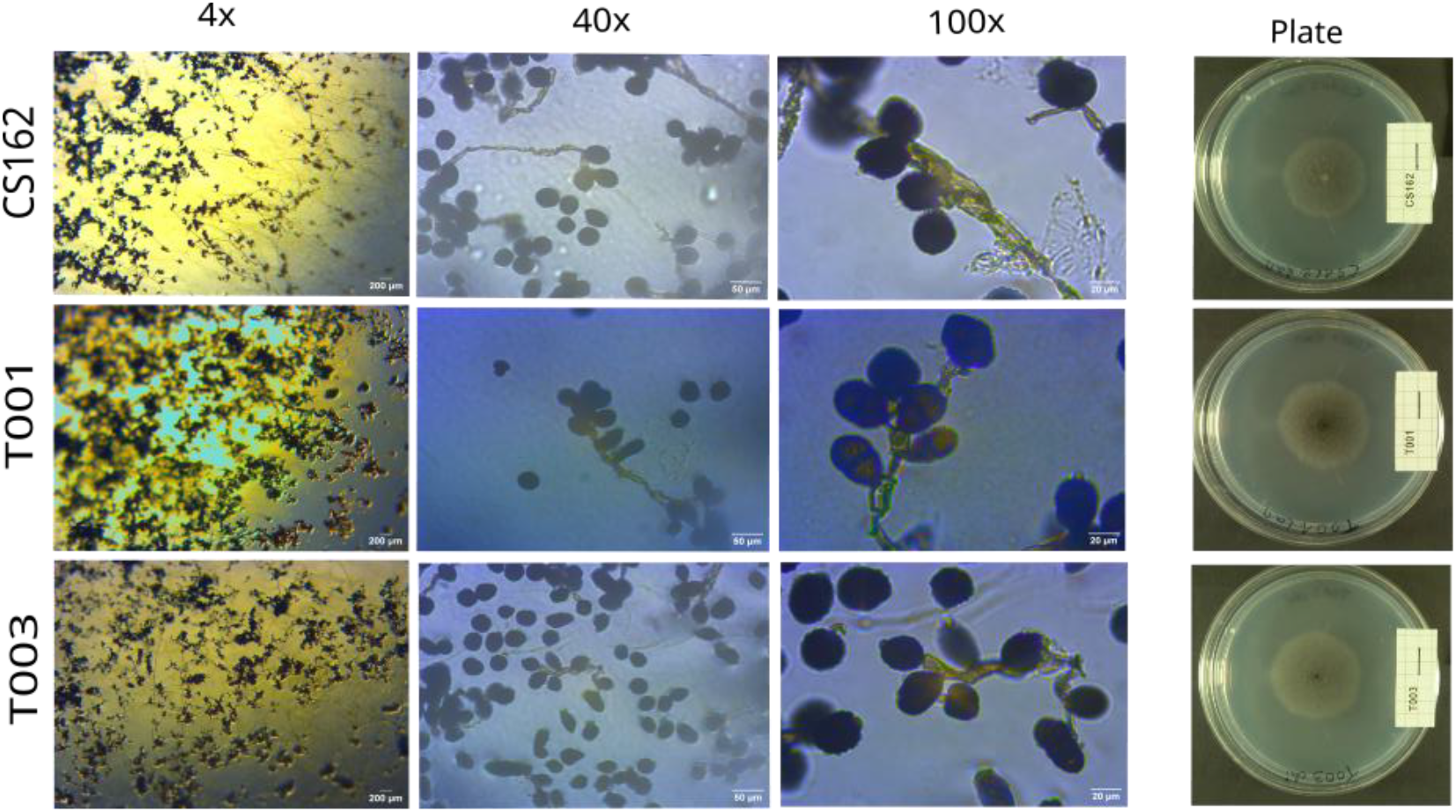
Conidia and colony morphology of *Alternaria atra* isolates CS162, T001, and T003. Light microscopy images were captured at 4×, 40×, and 100× magnification, showing typical conidial morphology. Colony morphology on SNA medium is shown on agar plates against a black background; plates were inoculated in the centre and photographed after 7 days of incubation.

Interestingly, whereas the conidial size, shape, and structure appear nearly identical for all three isolates, their colony morphology on agar plates differs visibly. All three isolates produced dark brown to black colonies on SNA medium. Isolates T001 and T003 showed a colony morphology comparable to *Alternaria alternata* CS046, which was reported as pathogen by (Schmey et al. 2023) and later used in this study as a positive control. In contrast, CS162 formed colonies of similar colour but with additional white, fluffy aerial hyphae on the surface, giving the centre of the colony a lighter appearance when viewed from above. This growth pattern is intrinsic to the fungus and is not due to contamination. A similar phenotype is observed when the fungus is grown on leaves (see below).

### Whole genome sequencing, assembly and annotation

The sequencing coverage reached 21× for T001 and 23× for T003, as reported by the assembler Flye. The newly assembled genomes for T001 and T003 comprised 177 and 139 contigs, respectively. Genome sizes were comparable between T001, T003, and CS162, whereas the downloaded genome MOD1FUNGI7 was notably smaller (table 1). BUSCO analyses indicated high completeness exceeding 98% for all four genomes (figure 4A). CS162 showed a higher proportion of duplicated BUSCOs compared to the other three genomes, while T001 and T003 exhibited genome sizes comparable to CS162 but duplication levels similar to those observed for MOD1FUNGI7. EarlGrey repeat annotation identified repeat content of 10.64% in T001 and 10.70% in T003, both higher than the repeat content observed in CS162 (8.15%), whereas MOD1FUNGI7 showed a lower proportion of annotated repeats at 3.73%.

**Table 1.** Assembly and annotation statistics. The assemblies T001 and T003 stem from this study, the assemblies CS162 and MOD1FUNGI7 are available from NCBI. Annotation followed a uniform funannotate protocol to ensure comparability.

| isolate | CS162 | T001 | T003 | MOD1FUNGI7 |
| --- | --- | --- | --- | --- |
| Assembly Size | 39,606,256 bp | 39,096,361 bp | 39,152,512 bp | 34,819,731 bp |
| Num Scaffolds | 43 | 177 | 139 | 927 |
| Percent GC | 50.44% | 50.88% | 50.87% | 50.97% |
| BUSCO scores | C:98.8%<br>[S:93.4%,<br>D:5.4%],<br>F:0.3%, M:0.9%<br>, n:6641, E:3.1% | C:98.8%<br>[S:98.2%,<br>D:0.7%],<br>F:0.3%, M:0.8%<br>, n:6641, E:3.2% | C:98.8%<br>[S:98.4%,<br>D:0.4%],<br>F:0.3%, M:0.9%<br>n:6641, E:3.2% | C:98.4%<br>[S:98.3%,<br>D:0.1%],<br>F:0.6%, M:1.0%<br>n:6641, E:3.1% |
| Repeat content | 8.15% | 10.64% | 10.70% | 3.73% |
| Num Genes | 12,472 | 12,247 | 12,122 | 11,678 |
| Num Proteins | 12,345 | 12,106 | 11,954 | 11,551 |
| Num tRNA | 127 | 141 | 168 | 127 |
| Unique Proteins | 386 | 407 | 336 | 356 |

### Mating types

In silico PCR with the first MAT1-1-1 primer pair produced products for CS162, T001, and T003. BLAST confirmed that these sequences correspond to the *Ulocladium atrum* (now *A. atra*) MAT1-1-1 locus. MOD1FUNGI7 did not yield a product. The MAT1-2-1 primer pair from the same study produced no products with three mismatches and generated only an abnormally long artifact for T003 when four mismatches were allowed. The second MAT1-1-1 primer pair did not produce any in silico PCR products for any of the four samples with either three or four mismatches. The MAT1-2-1 primer pair from Geng et al. generated a product for MOD1FUNGI7, which BLAST confirmed corresponds to the *Ulocladium atrum* (now *A. atra*) MAT1-2-1 locus.

The three isolates CS162, T001, and T003 showed high-identity BLASTn matches to the mating-type MAT1-1-1 sequence, which did not produce any match in the fourth isolate, MOD1FUNGI7. The MAT1-2-1 sequence only matched in MOD1FUNGI7 and not in the other three assemblies. As no assembly contained both idiomorphs, all four isolates are heterothallic, even though the query sequences originate from the same homothallic reference isolate.

### Phylogeny

Both genome-wide and marker-based analyses placed the studied isolates within the *Alternaria atra* clade together with reference sequences, providing strong support for their species assignment. The Mashtree cladogram based on k-mer clustering of whole-genome assemblies (figure 2A) resolved the broad relationships among all study and reference isolates, while the barcode-based phylogeny (figure 2B) provided finer resolution within the clade. Node support was high in both analyses for the placement of the two isolates in the clade with *A. atra*.

**Figure 2.**
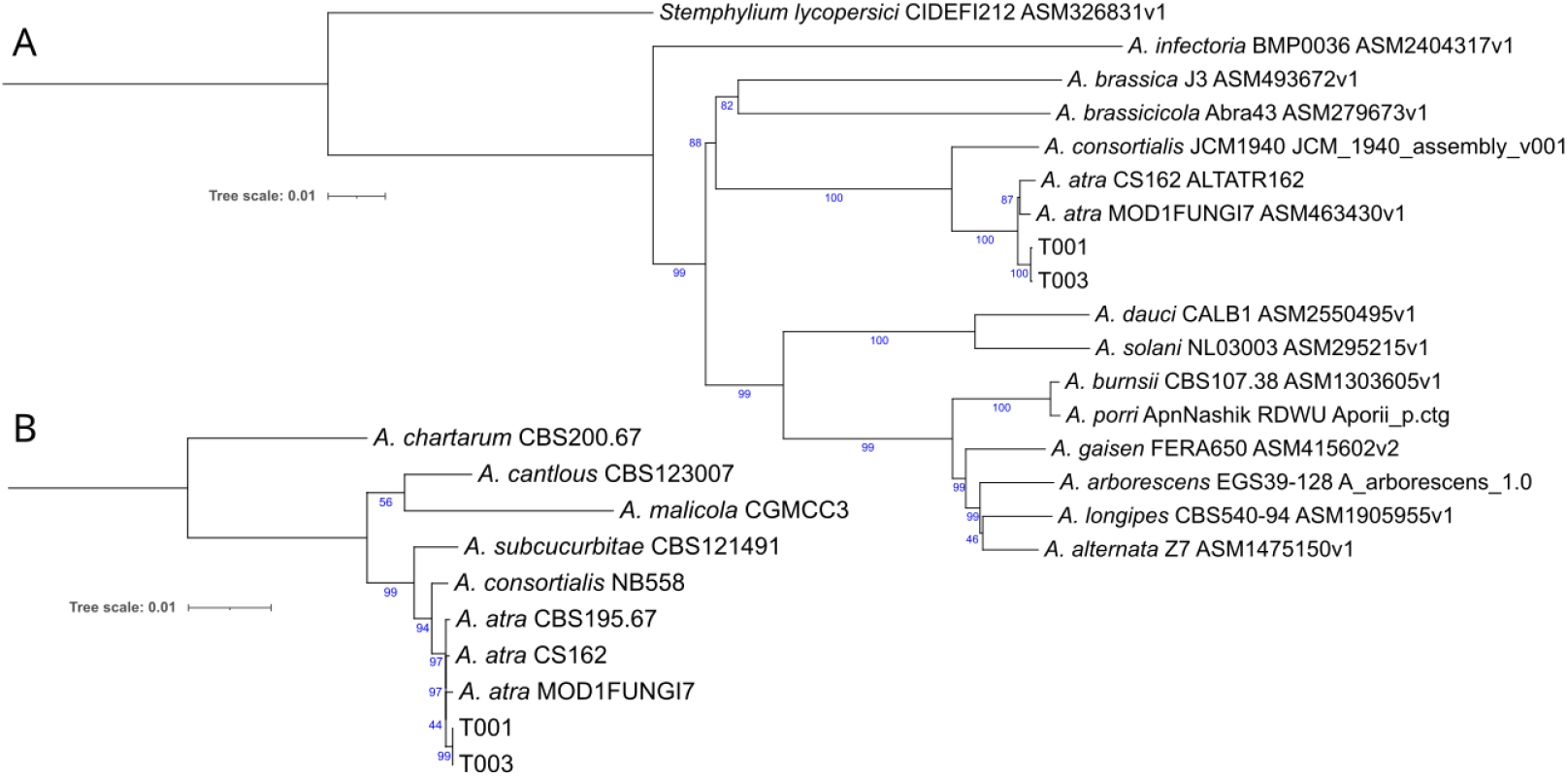
Genome-wide and marker-based relationships of study isolates and reference taxa. A) cladogram inferred from k-mer–based clustering of whole-genome assemblies using Mashtree. Species names are followed by isolate names and NCBI accession numbers. B) phylogeny reconstructed from concatenated barcode marker loci *alt a 1, gapdh, tef1*, and *rpb2*. Species names are followed by isolate names, the accession numbers of the sequences can be found in Gannibal et al. 2024. Node support values indicate bootstrap percentages based on 1000 replicates.

### Lifestyle under laboratory conditions

To assess host–isolate interactions experimentally, we performed drop inoculation assays on detached leaves of *Tillandsia landbeckii* (the host species of T001 and T003) and *Solanum chilense* (the host species of CS162) using conidial suspensions of all three isolates, alongside a positive control as reference for infection symptoms. After 6 dpi, nearly all leaves showed visible fungal growth, and by 10 dpi all inoculated leaves were clearly colonized (figure 3). All fungal isolates produced diffuse fungal growth that could be interpreted as necrotrophic infections. The mock-treated leaves of *S. chilense* (water control) remained free of fungal growth but showed humidity-associated stress symptoms. Overall fungal growth was fastest for the positive control *A. alternata* CS046, followed by the *A. atra* isolates T001, CS162, and T003. On *T. landbeckii*, fungal growth appeared mainly at the wound site where the leaf had been cut and inoculated. By 10 dpi, small amounts of fungal growth also developed on the leaf surfaces of *T. landbeckii* inoculated with CS046 and T001. While the positive control CS046 and the two *T. landbeckii* isolates T001 and T003 produced only dark colonies, CS162 additionally formed white aerial hyphae on both leaf types, consistent with the observations from SNA medium. The fungi isolated from the inoculated leaves matched those used for inoculation, thereby fulfilling Koch’s postulates.

**Figure 3.**
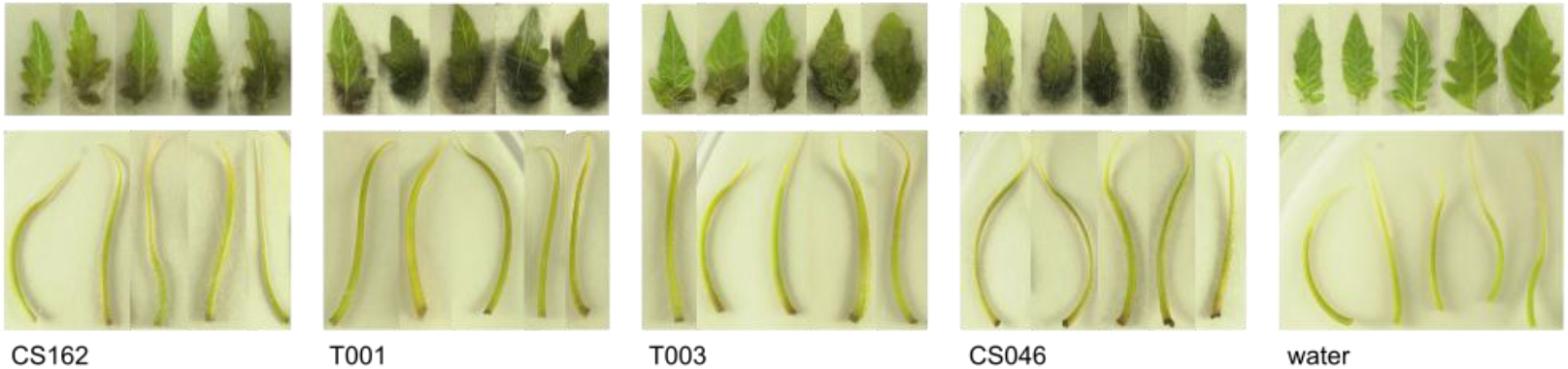
Detached leaf infection assays. Detached leaves of *Solanum chilense* and *Tillandsia landbeckii* inoculated with the *Alternaria atra* isolates, *Alternaria alternata* CS046 as pathogenic positive control, and mock-treated with water. Five representative leaves per treatment are shown. All inoculated fungi exhibited visible colonization phenotypes while there was no infection in the mock treatment.

### Comparative genomics of lifestyle indicators

Effector prediction revealed consistent levels of cytoplasmic effectors across all four isolates, whereas apoplastic effector abundances showed more pronounced variation (figure 4B). Across the genomes, between 26.7% and 29.0% of secreted proteins were predicted to be effectors. The proportion of cytoplasmic effectors was relatively stable, ranging narrowly from 6.0% to 7.1% of all secreted proteins, with absolute counts differing by only a few proteins between isolates. Apoplastic effector representation showed slightly more variance, spanning 20.7– 22.2% of secreted proteins.

**Figure 4.**
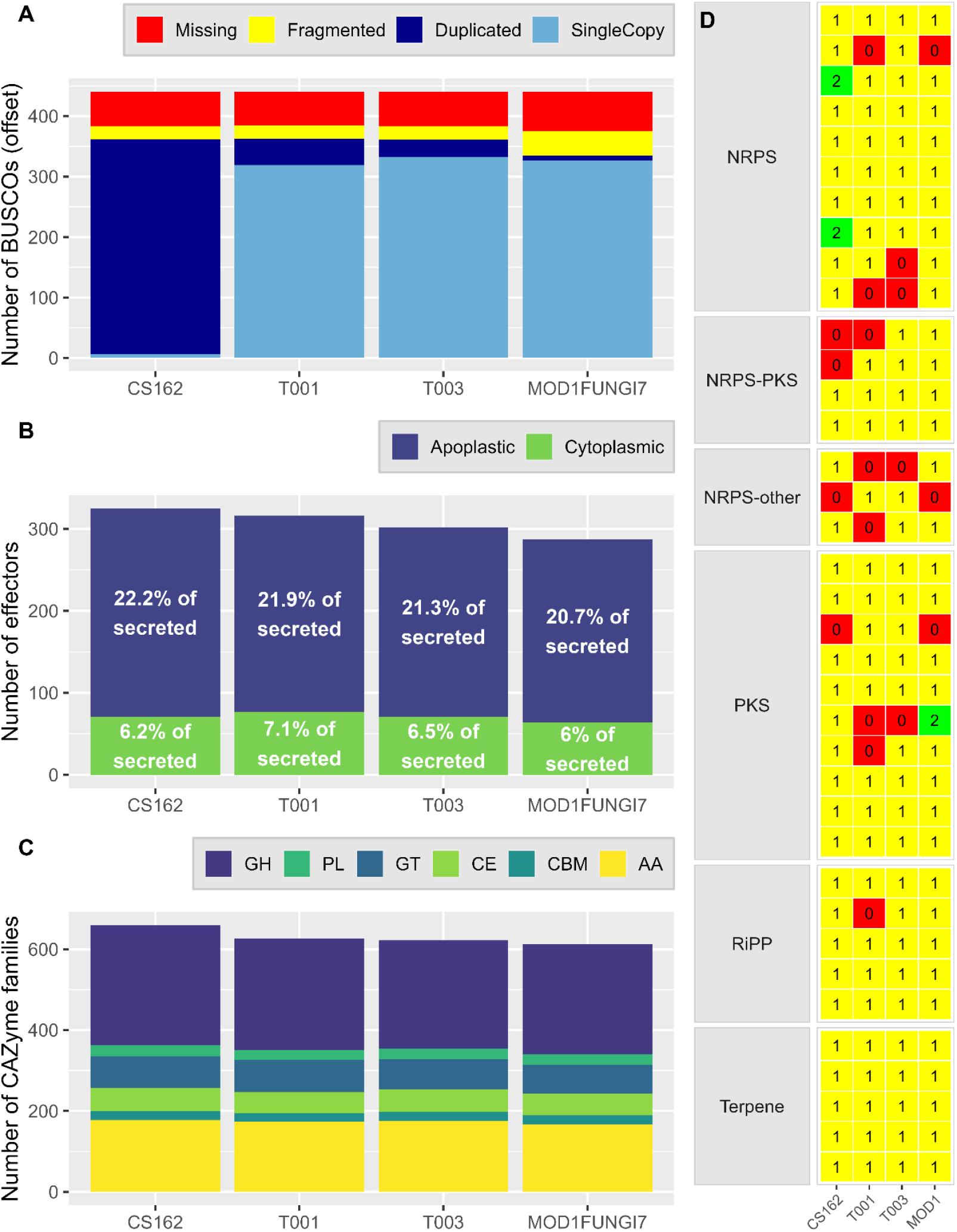
Comparative genomic features of the four fungal isolates. A) BUSCO completeness scores shown as stacked bar plots. To allow all categories to be visualized within a comparable scale, the values for single-copy BUSCOs were reduced by 6,200 in the plot. B) Predicted effector repertoires. Stacked bar plots show the absolute numbers of cytoplasmic and apoplastic effectors per isolate, and the percentages indicate how these counts relate to the total number of secreted proteins used as input for effector prediction. C) CAZyme family composition generated with funannotate compare, summarizing the major CAZyme classes present in each isolate. AA, auxiliary activities; CBM, carbohydrate-binding modules; CE, carbohydrate esterases; GH, glycoside hydrolases; GT, glycosyltransferases; PL, polysaccharide lyases. D) Presence–absence matrix of biosynthetic gene cluster (BGC) families identified with BiG-SCAPE v2.

We clustered the BGCs into gene cluster families (GCFs) with BiG-SCAPE. In the resulting presence-absence matrix (figure 4D), we aimed to identify GCFs associated with pathogenicity or necrotrophy by examining GCFs present in the presumed pathogen CS162 that are absent in both presumed endophytes T001 and T003. Only three GCFs followed this pattern: one unidentified NRPS, one unidentified NRPS-indole and the T1PKS for dehydrocurvularin. Consequently, we checked for GCFs that might be associated with plant beneficial functions present in T001 and T003 while absent in CS162, the presumed pathogen, and MOD1FUNGI7, the post-harvest rot. Only two GCFs were present in the presumed endophytes but absent in both others: one unidentified NRPS-indole and one unidentified PKS. Using a presence-absence matrix with such limited numbers of genomes does not provide the resolution needed to identify pathogenicity related or beneficial BGCs. As the two presumed endophytes share ecological niches, many genome features and all sequencing and assembly methods, the evaluation relied on these two genome annotations to be very similar. However, there are five GCFs present in only one of the two samples that were presumed similar.

Predicted CAZyme repertoires were broadly similar across the four *A. atra* isolates but showed quantitative differences among CAZyme families (figure 4C). Glycoside hydrolases (GH) represented the largest CAZyme class in all genomes and exhibited the greatest absolute variation among isolates, with higher counts in CS162 and lower counts in the endophytic isolates. Auxiliary activity enzymes (AA) were the second most abundant class and showed moderate variation, whereas glycosyltransferases (GT) and carbohydrate esterases (CE) differed only slightly among genomes. Carbohydrate-binding modules (CBM) and polysaccharide lyases (PL) were consistently present at lower and more uniform levels across all isolates. Although the CS162 assembly shows elevated BUSCO duplication levels (reported above), CAZyme family counts were not uniformly increased in this genome, and several categories displayed comparable abundance across isolates, indicating that duplication artifacts do not affect all CAZyme families equally. The CAZyme family AA9 comprises 34 members in all four genomes. The families GH5 and GH10 comprise 20 and 7 members, respectively, in three of the genomes, and 21 and 8 members, respectively, in CS162.

### Predictions of trophic modes using CATAStrophy

CATAStrophy predicted trophic lifestyles for the four genomes using three nomenclature frameworks, with consistent assignments under the traditional nomenclature and more variable outcomes under the alternative classification schemes (table 2). Under Nomenclature 1, all genomes were assigned necrotrophy as the closest centroid, while hemibiotrophy was consistently recovered among additional classes with high relative centroid distance (RCD) scores. Predictions under Nomenclature 2 differed among genomes, with vasculartrophy identified as the closest centroid for T001, T003, and CS162, and polymertrophy for MOD1FUNGI7; however, multiple alternative classes showed comparable RCD support across all genomes. Under Nomenclature 3, the closest centroid assignments also varied, with three genomes classified as polymertroph_narrow and one as mesotroph_intracellular, while overlapping alternative classifications were retained. The presence of several classes with similarly high RCD scores indicates that trophic classifications inferred from CAZyme profiles are not uniquely resolved for these genomes.

**Table 2.**
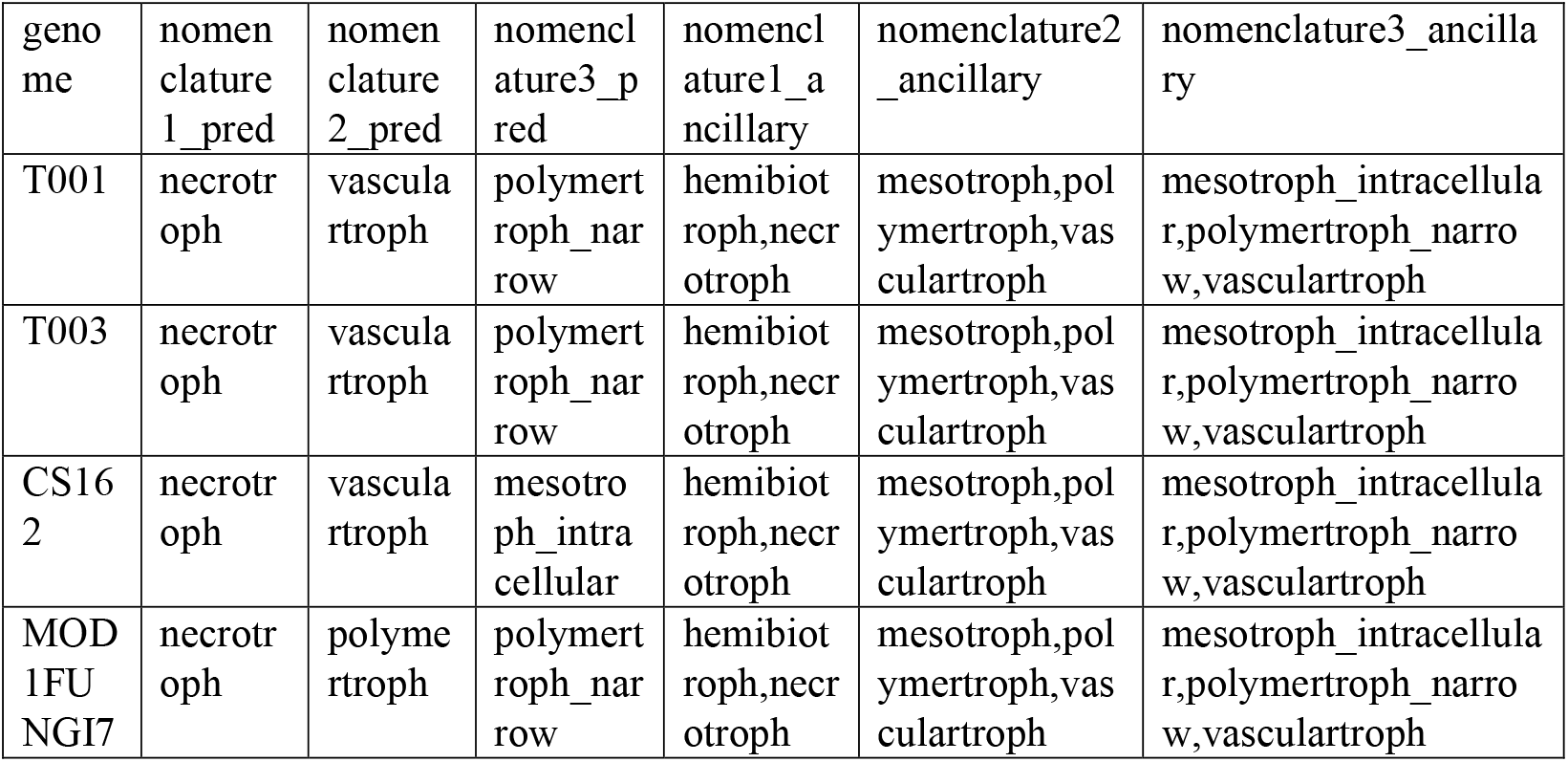
CATAStrophy results. Nomenclature 1 refers to traditional descriptions of fungal lifestyles, while Nomenclature 2 reflects the lifestyle classifications from the authors of CATAStrophy. Nomenclature 3 theoretically represents a refinement of Nomenclature 2. The first three columns are labelled pred and show the best prediction, determined by an RCD (relative centroid distance; see CATAStrophy documentation) score of 1. The columns labelled ancillary list all other predictions that had an RCD score above 0.8.

## Discussion

The lifestyle of *Alternaria* fungi has long been difficult to interpret because species boundaries within the genus have been obscured by extensive morphological plasticity and recurrent taxonomic revisions. In this study, all isolates were identified to species level using an integrative approach combining morphological characters and phylogenetic inference, allowing us to confidently assign them to *Alternaria atra* sensu lato. This is particularly important given the history of renaming and taxonomic confusion within *Alternaria*, where closely related taxa have frequently been synonymized or reclassified, complicating comparisons across studies. By demonstrating that the *A. atra* isolates analysed here form a well-supported, monophyletic group, our results confirm that the strains investigated belong to the same biological species, providing a robust framework for interpreting their shared genomic features and inferred lifestyle.

We present two draft *A. atra* genome assemblies that, despite being fragmented, are highly complete and represent valuable genomic resources. Both assemblies are comparable in size to the published CS162 genome, whereas MOD1FUNGI7 is substantially smaller. Genome size differences partly reflect biological variation in repeat content, which is slightly higher in the newly generated genomes than in CS162 and considerably lower in MOD1FUNGI7. The elevated duplicated BUSCO score observed in CS162 might potentially represent a technical artefact associated with reference-guided scaffolding. We consequently considered the impact of technical duplication artefacts on downstream analyses and assessed it to be minimal, as gene family sizes show both conserved and specifically expanded families rather than uniform inflation. Re-annotation of the genome CS162 using the funannotate pipeline yielded slightly improved results compared with the original publication (Bonthala et al. 2021), identifying 299 additional genes and 117 additional proteins.

Each genome contained only one mating type idiomorph: MAT1-1-1 in CS162, T001, and T003, and MAT1-2-1 in MOD1FUNGI7. This pattern supports the heterothallic system reported by (Gannibal and Gomzhina 2024) rather than the homothallic condition described by (Geng et al. 2014). Heterothallism may be viewed as being less consistent with a strictly endophytic lifestyle, as the likelihood of encountering a complementary mating type within plant tissues could be limited. Instead, a heterothallic reproductive mode is more consistent with at least occasional free-living or surface-associated phases, such as pathogenic growth on plant leaves or saprotrophic growth. Other endophytes, like *Pezicula neosporulosa*, show a mosaic of subpopulations with both heterothallic and homothallic strains (Yuan et al. 2022). The number of samples in our dataset is not sufficient to detect such patterns or to draw firm conclusions about the evolutionary or biological significance of mating-type variation in this context.

In the detached leaf assay, all three isolates were able to colonize the leaf tissue under laboratory conditions, despite differences in growth rate and apparent aggressiveness. Although collection data suggest that CS162 behaves as a pathogen and T001 and T003 as endophytes in nature, our results indicate that under standardized in vitro conditions, all isolates exhibit similar capacities for tissue invasion and growth. The strict surface sterilization procedure ensured that colonization observed in the assay reflects active fungal growth, supporting the idea that all three isolates possess the genetic and physiological potential required for tissue colonization. While growth on plates alone does not confirm a saprophytic lifestyle, the ability to develop on detached, possibly senescing leaf tissue suggests flexibility in nutrient acquisition strategies and is consistent with their potential to persist or grow as saprophytes under appropriate conditions.

Although the pathogenic genome CS162 shows a high number of duplicated BUSCOs, which may indicate potential assembly or annotation bias, this isolate nevertheless exhibits the highest number of predicted effectors. As effectors are important pathogenicity factors (Stergiopoulos and Wit 2009), this finding aligns with the general expectation. Comparative analyses across 84 plant-colonizing fungi highlighted that symbiotic fungi have a reduced effector repertoire compared to necrotrophic and saprophytic fungi (Lo Presti et al. 2015).

Scott et al. 2023 (Scott et al. 2023) reported that endophytic *Trichoderma* contained more BGCs than non-endophytes, likely reflecting the need for a wider range of metabolites to interact with host plants and other microbes. In contrast, in our study, the pathogen showed higher overall BGC counts than the other isolates, which could reflect adaptations for plant infection or technical assembly artefacts. We detected more differences between the two endophytes than between the pathogen and the endophytes, making claims about potential pathogenicity factors and endophytism markers unreliable.

Previous studies have identified specific CAZyme families as being associated with an endophytic lifestyle. Yuan et al. reported an enrichment of GH5 and AA9 family members in plant-beneficial endophytic fungi relative to pathogenic relatives (Yuan et al. 2022), while Mesny et al. identified GH10 and AA9 as part of a conserved gene repertoire linked to endophytism (Mesny et al. 2021). We observed highly similar copy numbers for all three CAZyme families, with identical numbers for AA9 in all four genomes. The families GH5 and GH10 had similar numbers in the two presumed endophytes and the downloaded genome of the post-harvest rot, but one additional member in the presumed pathogen. Given that the presumed pathogen had generally more genes annotated, this may reflect technical bias and the CAZyme repertoires appear conserved. Rather than indicating a lack of lifestyle differentiation, this conservation may reflect the capacity of these isolates to adopt pathogenic, endophytic, or postharvest lifestyles depending on host context or environmental conditions. Under this scenario, CAZyme repertoires may represent a shared genomic toolkit that supports lifestyle plasticity.

CAZyme-based trophic predictions consistently suggest necrotrophic tendencies under traditional classification, but assignments were more variable under alternative frameworks, reflecting ambiguity in fine-scale trophic resolution. These results indicate that while our isolates likely employ broadly necrotrophic or polymer-degrading strategies, their exact lifestyles may be flexible or intermediate.

Previous comparative genomic studies have similarly shown that CAZyme repertoires can overlap across necrotrophic, hemibiotrophic, saprotrophic, and endophytic Ascomycetes, with necrotrophs and hemibiotrophs often having expanded suites of cell-wall-degrading enzymes compared to biotrophs, and endophytes and generalist plant-associated fungi exhibiting intermediate or mixed CAZyme and effector profiles that make clear lifestyle delimitation difficult from enzyme content alone (Hill et al. 2022; Wang et al. 2022; Lyu et al. 2015).

In conclusion, our findings support lifestyle plasticity in *Alternaria atra*, as isolates with different collection histories exhibit similar behaviour on different plant hosts under laboratory conditions and lack genomic features indicative of a specific ecological lifestyle. The two newly generated genome assemblies provide valuable resources that expand available genomic data for this species and enable future investigations into regulatory and environmental drivers of fungal ecological diversity.

## Data availability

The ONT sequencing raw reads and the generated assemblies for T001 and T003 are deposited in the European Nucleotide Archive (ENA) project PRJEB105036. Annotation data of the newly generated genomes and re-annotation data of the publicly available genomes are deposited on zenodo under https://doi.org/10.5281/zenodo.18785799.

## Acknowledgements

We thank Stefan Krebs for ONT sequencing. We also thank Maie Bachmann and Bettina Bastian for help with laboratory experiments.

